# DNA replication protein Cdc45 directly interacts with PCNA via its PIP box in *Leishmania donovani* and the Cdc45 PIP box is essential for cell survival

**DOI:** 10.1101/831669

**Authors:** Aarti Yadav, Varshni Sharma, Jyoti Pal, Pallavi Gulati, Manisha Goel, Udita Chandra, Neha Bansal, Swati Saha

## Abstract

DNA replication protein **C**dc45 is an integral part of the eukaryotic replicative helicase whose other components are the **M**cm2-7 core, and **G**INS. We identified a PIP box motif in *Leishmania donovani* Cdc45. This motif is typically linked to interaction with the eukaryotic clamp **p**roliferating **c**ell **n**uclear **a**ntigen (PCNA). The homotrimeric PCNA can potentially bind upto three different proteins simultaneously via a loop region present in each monomer. Multiple binding partners have been identified from among the replication machinery in other eukaryotes, and the concerted /sequential binding of these partners are central to the fidelity of the replication process. Though conserved in Cdc45 across *Leishmania* species and *Trypanosoma cruzi,* the PIP box is absent in *Trypanosoma brucei* Cdc45. Here we investigate the possibility of Cdc45-PCNA interaction and the role of such an interaction in the *in vivo* context. Having confirmed the importance of Cdc45 in *Leishmania* DNA replication we establish that Cdc45 and PCNA interact stably in whole cell extracts, interacting with each other directly *in vitro* also. The interaction is mediated via the Cdc45 PIP box. This PIP box is essential for *Leishmania* survival. The importance of the Cdc45 PIP box is also examined in *Schizosaccharomyces pombe,* and it is found to be essential for cell survival in this organism also. Our results implicate a role for the *Leishmania* Cdc45 PIP box in recruiting or stabilizing PCNA on chromatin. The Cdc45-PCNA interaction might help tether PCNA and associated replicative DNA polymerase to the DNA template, thus facilitating replication fork elongation. Though multiple replication proteins have been identified to associate with PCNA in other eukaryotes, this is the first report demonstrating a direct interaction between Cdc45 and PCNA, and while our analysis suggests the interaction may not occur in human cells, it indicates that it is not confined to trypanosomatids.

**Author Summary:** Leishmaniases are manifested in three forms: cutaneous, sub-cutaneous and visceral. The prevalent form in the Indian subcontinent is visceral Leishmaniasis (VL), which is fatal if not treated on time. While there are drugs for treatment, the hunt for additional drugs continues due to emerging drug resistance patterns. The parasite is transmitted by the bite of the sandfly, whereupon it establishes itself within cells of the host immune system (macrophages) and reproduces by binary fission. The replication of its genome is essential for parasite survival. Eukaryotic DNA replication is generally conserved across species. This study targets Cdc45, a protein that helps unwind the DNA double helix to enable copying of the two strands into two daughter strands. The new chains of DNA are synthesized by DNA polymerases, and a trimeric protein, proliferating cell nuclear antigen (PCNA), helps clamp the polymerases onto the template. In this study we find Cdc45 to interact with PCNA, and have identified the motif in Cdc45 via which it does so. Our results suggest this interaction is seen in some other eukaryotes as well. Based on the results of our experiments we propose that Cdc45 may help moor PCNA-polymerase complexes to template DNA.

## Introduction

Leishmaniases afflict 12 million people across 88 countries, and are mainly manifested as cutaneous, sub-cutaneous and visceral leishmaniasis. Visceral Leishmaniasis remains a threat due to emerging drug resistance as well as due to risks associated with dual HIV-*Leishmania* infections, and is deadly if not treated in a timely manner. In the Indian subcontinent visceral Leishmaniasis is caused by *Leishmania donovani*. The parasite is digenetic, shuttling between the insect host (sandfly) and the mammalian host. It reproduces asexually by binary fission in both hosts and DNA replication is central to the process.

The conserved process of eukaryotic DNA replication is marked by the licensing of origins in late mitosis/G1 followed by the firing of these origins in S phase. The licensing of origins is coupled to the formation of pre-replication complexes (pre-RCs) at or very near origins (reviewed in [1, 2]). The assembly of pre-RCs begins with the binding of the origin recognition complex ORC (comprising Orc1-6) to DNA, followed by the sequential recruitment of Cdc6, and two copies of Cdt1-MCM (MCM: Mcm2-7 heterohexameric complex). The association of the MCM double hexamer “licenses” origins to fire. As cells enter S phase Mcm4 and Mcm6 are phosphorylated by Dbf4-dependent kinase (DDK), leading ultimately to the association of Cdc45 and the heterotetrameric GINS (**g**o-**i**chi-**n**i-**s**an) along with other factors like Mcm10 [3, 4], that help convert the inactive double hexamer MCM complex into the active replicative helicase comprising Cdc45, a single MCM hexamer and GINS: the CMG complex (**C**dc45-**M**cm2-7-**G**INS; [5, 6]). Thus, two CMG complexes are formed at each licensed origin, and when the origin fires they move in opposite directions. EM and cross-linking studies with *Drosophila* CMG coupled with crystal structure analysis of human Cdc45 reveal that Cdc45 interacts with Mcm2-7 at the Mcm2/5 interface, while GINS interacts with Mcm2-7 at the Mcm3/5 interface. Cdc45 and GINS also interact with each other in this complex. As the Mcm2/5 interface forms the gate of the annular Mcm2-7 hexamer that closes once the Mcm2-7 ring has encircled the DNA template, it is concluded that Cdc45 safeguards the closed gate, thus ensuring the template DNA remains within the ring to enable translocation of the helicase during the elongation phase of replication [7–9]. This forms the basis for the activation of MCM helicase activity by Cdc45-GINS. The replisome that advances with the replication fork comprises a host of proteins including the CMG complex but not the ORC, with data from EM studies identifying a core replisome consisting of 20 subunits [10–13].

With orthologs of several of the conserved eukaryotic replication proteins being identified in trypanosomatids, DNA replication in trypanosomes seems to broadly resemble that of other eukaryotes. However, that trypanosomatids diverged from other eukaryotes very early on is reflected in the fact that some of the conserved replication proteins are absent [14–16]. While most of the components of the replication elongation machinery including Mcm2-7, Cdc45, GINS, the DNA polymerases, and PCNA are present [14,17–21], the same is not the case with components of the pre-replication complex. Thus, while Orc1 and Orc4 are present, Orcs 2, 3, 5 and 6 have not been found, neither has Cdt1 [22–24]. Furthermore, there is evidence of a divergent origin recognition complex (ORC) in *T. brucei* that comprises Orc1, Orc4, Orc1B (an Orc1-like protein), and two other proteins Tb7980 and Tb3120 that share very little sequence homology with the conserved Orc proteins [14, 19]. The *Trypanosoma brucei* CMG complex has been found to exhibit helicase activity *in vitro*, and knockdown of individual components of the *T. brucei* CMG complex results in defects in DNA replication and cell proliferation [19].

As the Cdc45 protein is highly conserved in sequence among trypanosomatids the *Leishmania* Cdc45 is expected to behave similarly. However, sequence analysis revealed that the *Leishmania* Cdc45 protein possesses a PIP box, a motif that is usually involved in interactions with proliferating cell nuclear antigen (PCNA). This motif is not evident in *Trypanosoma brucei* Cdc45. The present study was undertaken with the aim of addressing the functional role of this PIP motif, if any, in relation to DNA replication and cell survival.

## Results

### Depletion of Leishmania donovani Cdc45 leads to severe growth, cell cycle and DNA replication defects

The *Leishmania donovani* 1S *cdc45* gene was cloned as described in Supplementary Methods, and the gene sequenced (GenBank Accession no. MN612783). Previous studies in *T. brucei* have shown that Cdc45 is nuclear in G1 and S phase but is cytosolic after G2, and it was proposed that this nuclear export contributed to the prevention of DNA re-replication in the same cell cycle [19]. As Clustal Omega analysis [25] revealed *Leishmania donovani* Cdc45 to share ∼ 43% identity and ∼ 60% similarity with Cdc45 from *T. brucei* and *T. cruzi* (S1 Fig) we initiated our study with examining the possibility of a similar mechanism of replication regulation existing in *Leishmania*. The gene encoding LdCdc45 lies on chromosome 33, and in the aneuploid *Leishmania donovani* 1S genome this chromosome is trisomic [26]. To analyze subcellular localization of the Cdc45 protein we tagged one of the genomic alleles with *eGFP* using homologous recombination as described in Supplementary Methods, and verified the authenticity of the recombinants by PCRs across the replacement junctions (S2a Fig). After confirming expression of full length Cdc45-eGFP by western blot analysis (S2b Fig), Cdc45-eGFP expression within these cells was examined by indirect immunofluorescence with anti-eGFP antibodies, using kinetoplast morphology and segregation pattern as cell cycle stage indicator [27]. LdCdc45 behaved differently from TbCdc45, remaining nuclear throughout the cell cycle and thus ruling out the possibility of nuclear export being a mode of replication regulation (S2c Fig).

The effects of Cdc45 depletion on cell growth and cell cycle progression were investigated by creating genomic knockouts as *Leishmania* cells lack canonical RNAi machinery. The process of making knockout lines was complicated by the fact that there must be three genomic alleles as the gene lies on a trisomic chromosome. Accordingly, we attempted to sequentially knock out all three alleles by homologous recombination, as described in Supplementary Methods. In the first round of homologous recombination we created two individual heterozygous knockout lines where one allele was replaced by either the hygromycin resistance cassette or the neomycin resistance cassette (*cdc45*^-/+/+^ ::hyg and *cdc45*^-/+/+^ ::neo respectively; Figs. 1a and 1b). A second allele in the *cdc45*^-/+/+^ ::hyg line was next replaced with the neomycin resistance cassette to create a double heterozygous knockout line *cdc45*^-/-/+^ (Fig. 1c). While all attempts to create a true *cdc45*-null failed we were able to successfully create a *cdc45*-null in a Cdc45^+^ background (Cdc45-FLAG expressed ectopically from a plasmid in *cdc45*^-/-/+^cells, described in Supplementary Methods; Fig. 1d). All the lines were authenticated by PCRs across the deletion junctions at both ends (Figs. 1a-1d).

**Fig. 1:**
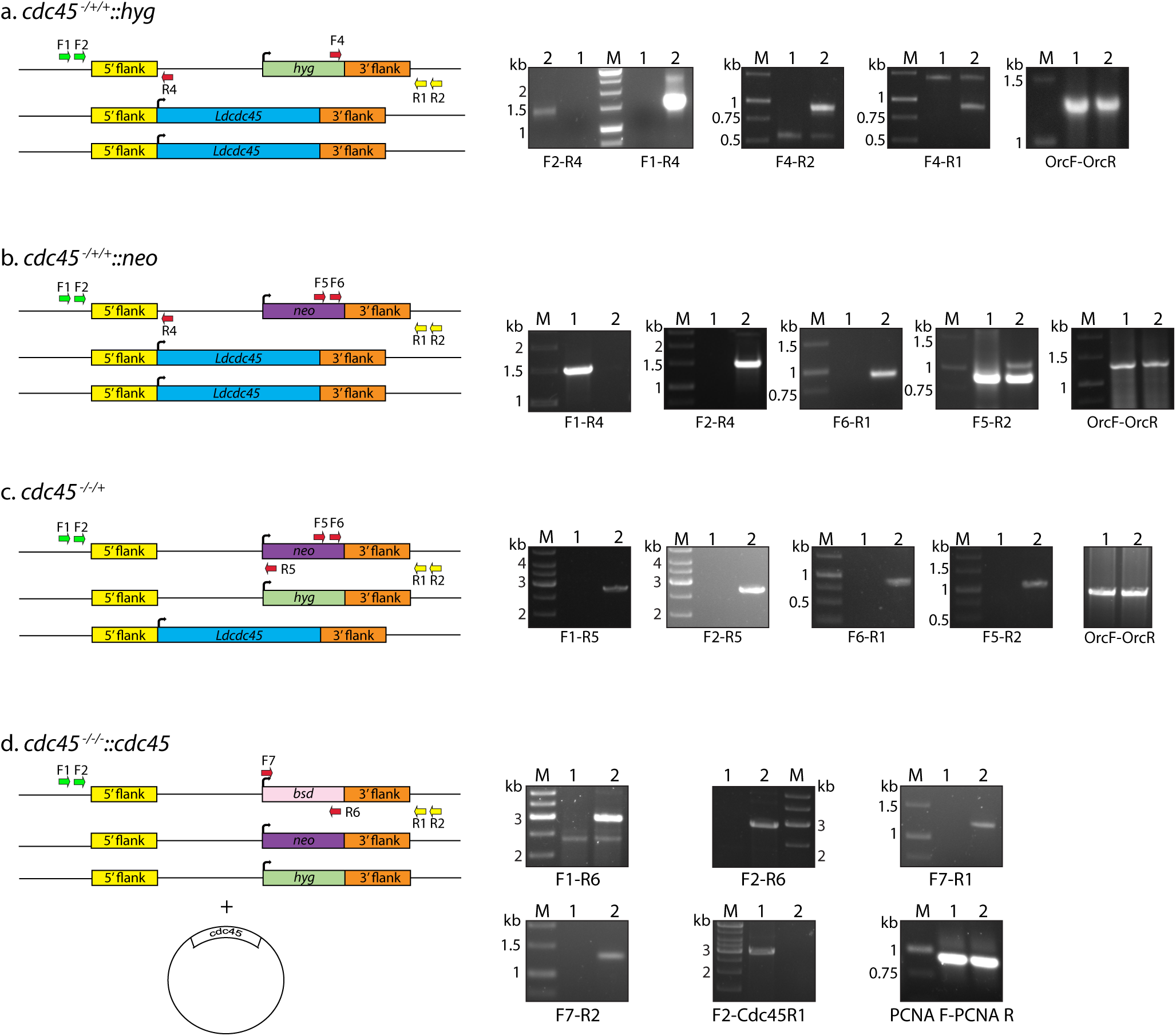
Creation of *cdc45* genomic knockouts: **a:** Replacement of one genomic allele with *hyg^r^* cassette. **b.** Replacement of one genomic allele with *neo^r^* cassette. **c.** Replacement of second genomic allele in *cdc45^-/-/+^::hyg* with *neo^r^* cassette. **d.** Replacement of third genomic allele in *cdc45^-/-/+^::cdc45* strain with *bsd^r^* cassette. Line diagrams represent schematics of knockout lines created. Primers used in screening indicated in line diagrams. Agarose gels depict screening of knockout lines. Primer pairs used in each case are indicated below the gel. Lanes 1-Ld1S, lanes 2-respective knockout line, M-DNA ladder marker. OrcF-OrcR and PCNAF-PCNAR PCRs served as positive controls.

The extent of expression of *cdc45* in *cdc45*^-/+/+^::hyg and *cdc45*^-/-/+^ cells was determined by real time PCR analyses of RNA isolated from logarithmically growing promastigotes. When *cdc45* expression in the knockout cells relative to in wild type cells was quantitated using the 2^-ΔΔC^_T_ method [28] it was found that while expression in *cdc45*^-/+/+^ ::hyg cells was ∼ 0.7 fold that seen in wild type cells, expression in *cdc45*^-/-/+^ cells was only ∼ 0.2 fold that seen in wild type cells (Fig. 2a) - suggesting that the three alleles may not be equivalently expressed. Analysis of the growth patterns of the single allele knockouts *cdc45*^-/+/+^::hyg and *cdc45*^-/+/+^ ::neo revealed that both these lines grew similarly, growing slower than control cells and never reaching the same cell density as controls, but reaching stationary phase at around the same time as control cells (Fig. 2b left panel). The double allele knockout *cdc45*^-/-/+^ cells grew even slower, reaching stationary phase a day later (Fig. 2b right panel). The slower growth of *cdc45*^-/-/+^ cells was linked to an increase in generation time to ∼16.55 h as compared to ∼9.43 h in case of control cells (Fig. 2c).

**Fig. 2:**
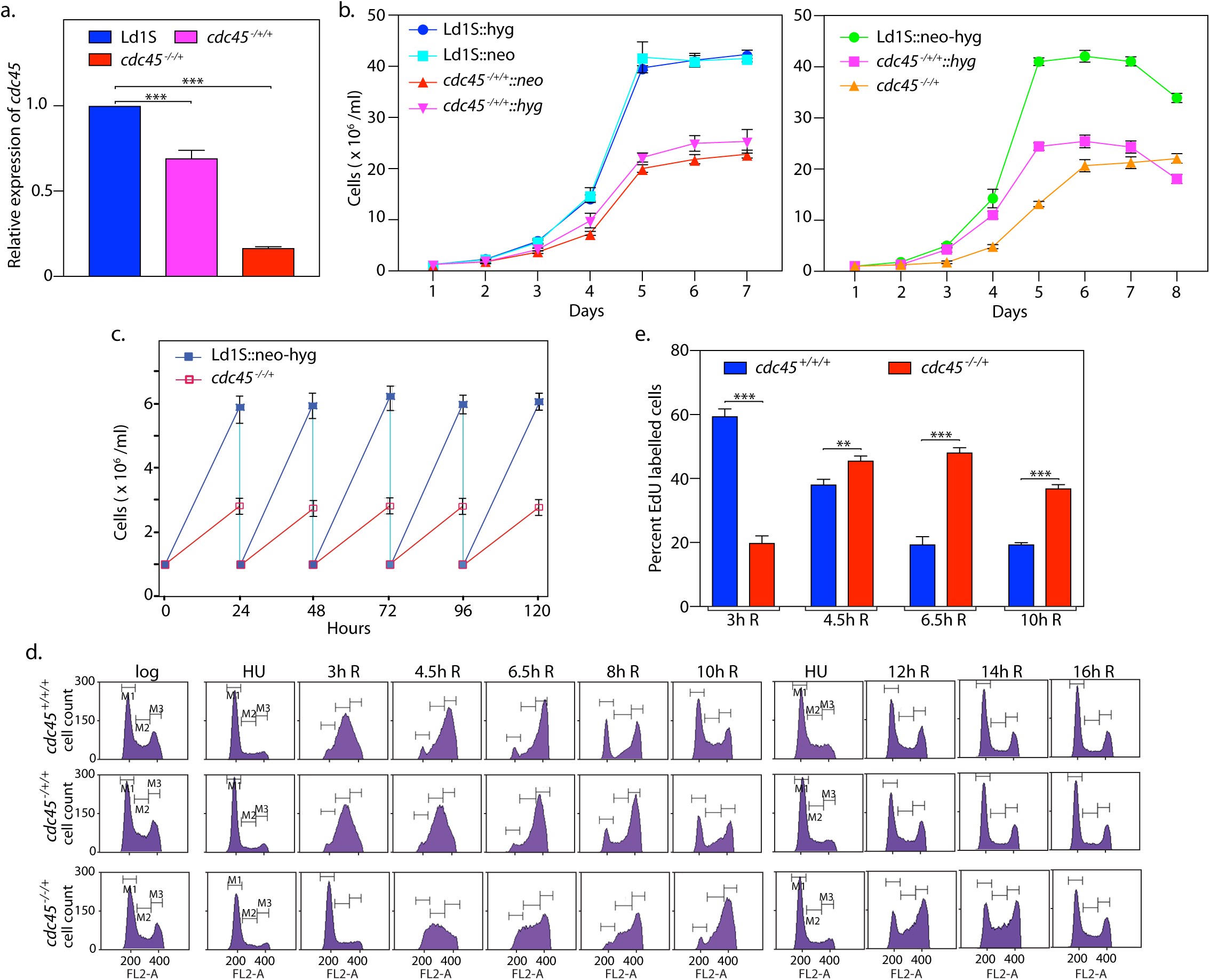
Depletion of Cdc45 causes growth and cell cycle aberrations. **a.** Analysis of *cdc45* expression in genomic knockout lines using real time PCR analysis. Tubulin served as internal control. The average of data from three experiments is presented here. Error bars represent standard deviation. Two-tailed student’s t-test was applied: *** *p* < 0.0005. **b.** Analysis of growth of Cdc45-depleted cells. Growth initiated from stationary phase cells, at 1×10^6^ cells/ml. Left panel: single allele replacement lines compared to control cells. Right panel: double allele and single allele replacement lines compared to control cells. In each case three experiments were set up in parallel and the graphs represent average values, with error bars representing standard deviation. **c.** Determination of generation time of double allele replacement cells *cdc45^-/-/+^* in comparison with control cells. Growth initiated at 1×10^6^ cells/ml from exponentially growing promastigotes, and cells diluted to 1×10^6^ cells/ml every 24 hours. Average of three experiments plotted, and error bars depict standard deviation. **d.** Flow cytometry analysis of Cdc45-depleted cells in comparison with control cells. Cells were synchronized at G1/S by HU treatment and then released into S phase. Time-points at which cells were sampled are indicated above each column of histograms *e.g.* 3h R signifies 3 hours after release from HU. M1, M2 and M3 gates represent cells in G1, S and G2/M respectively. 30000 events were counted in each case. The experiment was done thrice, and one data set is presented here. **e.** HU-treated cells were released into S phase and 1 ml aliquots pulsed with EdU 3 hours, 4.5 hours, 6.5 hours and 10 hours after release (15 min pulses). Data presented in bar chart are average of three experiments and error bars represent standard deviation. Two-tailed student’s t-test was applied: \*\**p* < 0.005; \*\*\**p* < 0.0005.

To assess the effect of Cdc45 depletion on cell cycle progression we synchronized cells with hydroxyurea at G1/S, released the cells into S phase, and monitored their progress across S and G2/M and thereafter back into G1 by flow cytometry analysis at various time-points. The patterns observed were distinctive (Fig. 2d). *cdc45*^-/+/+^ ::hyg cells entered S phase smoothly and reached mid-S phase at rates comparable to control cells, thereafter slowing down and beginning to re-enter G1 ∼ two hours later than control cells. *cdc45*^-/-/+^ cells showed a more acute phenotype, with cells entering and reaching mid-S phase later than usual, in general navigating S and G2/M phases much slower, and beginning to re-enter G1 phase ∼ four hours later than control cells. Analysis of *cdc45*^-/-/+^ cells by pulse-labeling with EdU at different time-points after release from HU-induced block and evaluating the EdU uptake (described in Methods), confirmed that depletion of Cdc45 leads to aberrant DNA replication patterns, with the process of DNA synthesis being more protracted (Fig. 2e).

### Defective phenotypes associated with Cdc45 depletion are rescued by ectopic expression of Cdc45

To confirm that the defective phenotypes observed were due to Cdc45 depletion we expressed Cdc45-FLAG in *cdc45*^-/-/+^ cells ectopically (described in Supplementary Methods; western blot analysis of whole cell extracts of transfectant cells in Fig. 3a) and analyzed the growth and cell cycle patterns of these cells. We found that episomal expression of Cdc45-FLAG in *cdc45*^-/-/+^ cells permitted the cells to grow at rates similar to wild type cells (Fig. 3b), and promoted normal cell cycle progression, with cells traversing S phase and G2/M and re-entering G1 phase in timely manner (Fig. 3c). These data lead us to conclude that the phenotypes of *cdc45*^-/-/+^ cells are due to Cdc45 depletion.

**Fig. 3:**
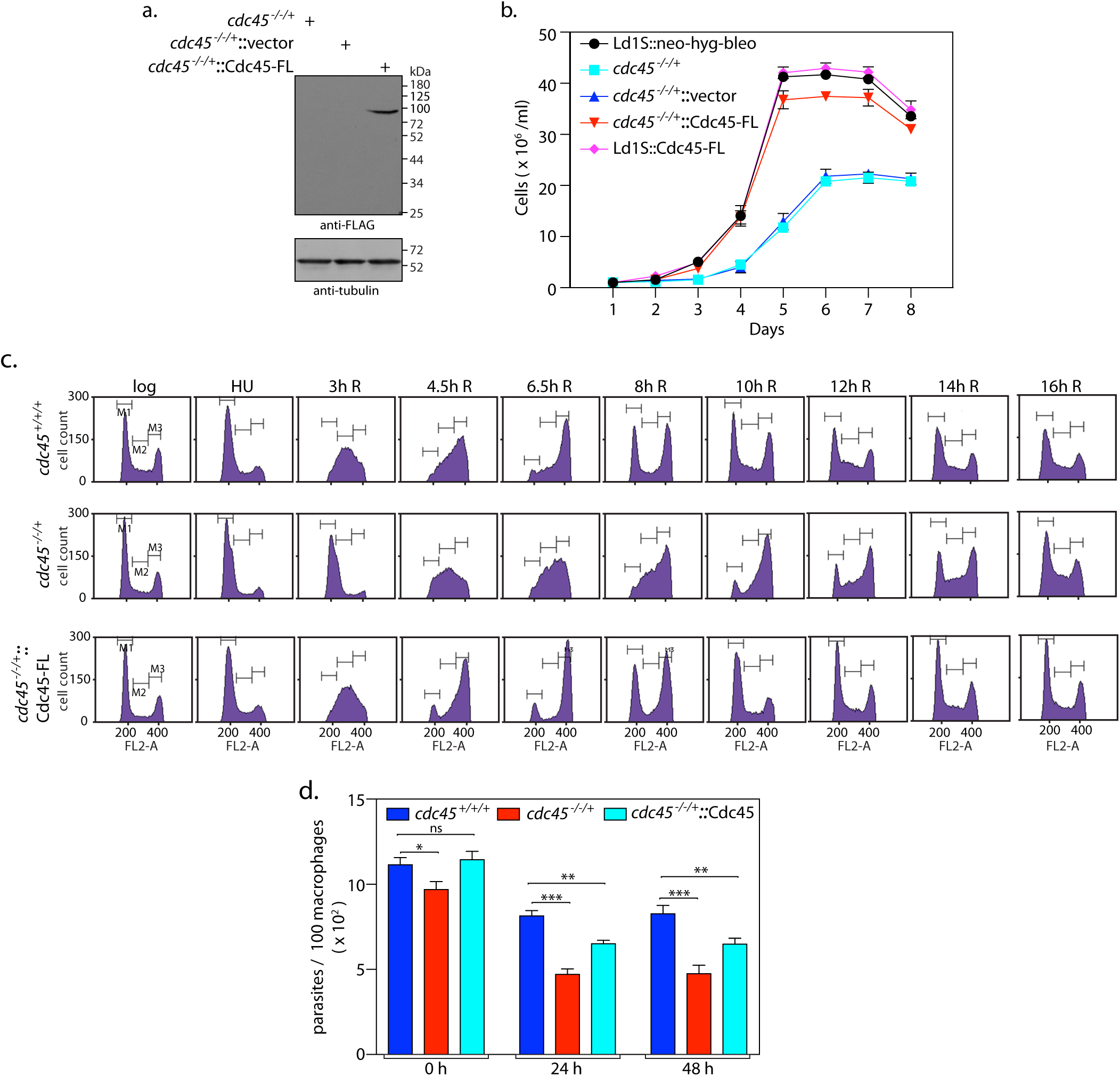
Ectopic expression of Cdc45 in *cdc45^-/-/+^* cells allows the cells to overcome defects associated with Cdc45 depletion. **a.** Western blot analysis of cell extracts using anti-FLAG antibody (Sigma, 1:1000 dil). *cdc45*^-/-/+^/vector: *cdc45*^-/-/+^ harboring pXG-FLAG(bleo) vector. *cdc45*^-/-/+^/Cdc45-FL: *cdc45*^-/-/+^ harboring pXG-Cdc45-FLAG(bleo). Loading control: tubulin. **b.** Analysis of growth pattern of rescue line in comparison with *cdc45^-/-/+^*cells and control cells. Growth initiated at 1×10^6^ cells/ml from stationary phase cells. Graphs represent average values of three experiments, with error bars representing standard deviation. **c.** Flow cytometry analysis of rescue line in comparison with *cdc45^-/-/+^*cells and control cells. 30000 events counted in every sampling. Sampling times indicated above the histograms. Cells representing G1, S and G2/M phases indicated by gates M1, M2, M3. Experiment done thrice, one data set depicted here. **d.** Analysis of cell survival within host macrophages (the parasites exist in insect host as non-infective procyclic promastigotes which later get differentiated into infective metacyclic promastigotes. The metacyclics are released into the mammalian host bloodstream upon insect bite where they establish residence in macrophages and multiply by binary fission). Parasites were scored by Z-stack imaging of DAPI-stained infected macrophages using confocal microscopy. Bar chart depicts averages of three experiments; error bars indicate standard deviation. Two-tailed student’s t-test was applied: \*\**p<*0.005; \*\*\**p* < 0.0005; ns-not significant.

Survival of the *cdc45*^-/-/+^ parasites within host macrophages was examined by infecting macrophages with metacyclic parasites, and analyzing the number of intracellular parasites right after infection as well as 24 hours and 48 hours later. We observed that while *cdc45*^-/-/+^ promastigotes were able to infect macrophages to similar extent as control parasites, they were subsequently unable to propagate to the same extent. The ectopic expression of Cdc45-FLAG in *cdc45*^-/-/+^ parasites partially rescued these defects (Fig. 3d). The reasons for a partial rescue are not clear; it is possible that the levels/extent of ectopic expression in the intracellular parasites varies from one to the other, leading to variable propagation rates.

### Analysis of LdCdc45 structure

Multiple sequence alignment of LdCdc45 with Cdc45 of other eukaryotes revealed that LdCdc45 shares 18-23% identity and 30-40% similarity with Cdc45 of the model eukaryotes (S3 Fig). Additionally, LdCdc45 harbors two exclusive stretches of ∼ 50 and 40 amino acid residues near the C-terminal end. When analyzing the amino acid sequence of LdCdc45 the presence of a PIP box (QRKLVEF) was detected between residues 500-506. This PIP box was conserved in all *Leishmania* species and was found in *T. cruzi* also, but not in *T. brucei* (S1 Fig). The PIP (**<u>P</u>**CNA-**i**nteracting **<u>p</u>**eptide) box is a short motif: **Q** x x **L/V/I** x x **F/Y F/Y** (where x is any amino acid) found in several proteins that interact with proliferating cell nuclear antigen (PCNA). PCNA, the eukaryotic ortholog of the bacterial β-clamp protein, is a homotrimeric protein forming an annular structure with pseudo-hexameric symmetry that enables it to encircle the DNA template and slide along. The PCNA monomer consists of two globular domains connected by a flexible **i**nter**<u>d</u>**omain **<u>c</u>**onnector **l**oop (IDCL). These monomers are arranged in such a way that the inner cavity of the trimeric ring has a positively charged surface comprising of α helices for association with DNA, while the outer surface of the ring comprises largely of β sheets. As it tethers the eukaryotic DNA polymerases to the DNA template it increases their processivity, and thus it plays a major role in both, DNA replication as well as DNA repair. Though not yet identified to interact with Cdc45 in any species, PCNA interacts with multiple other components of the DNA replication and DNA repair machinery (reviewed in [29]). With the identification of a PIP motif in LdCdc45, we analyzed the sequences of Cdc45 proteins of other eukaryotes. Cdc45 of *Schizosaccharomyces pombe, Drosophila melanogaster* and *Homo sapiens* were found to harbor PIP motifs, but not *Saccharomyces cerevisiae* Cdc45 (S3 Fig).

The CMG complex is an eleven subunit complex comprising Cdc45, Mcm2-7 heterohexamer and GINS heterotetramer (subunits: Sld5, Psf1, Psf2 and Psf3). The structure of the CMG complex was deciphered by single particle electron microscopy (EM) studies carried out using the recombinant *Drosophila* proteins [7], which revealed Cdc45 and GINS to both interact with Mcm2-7 on one side of the heterohexamer, with Cdc45 and GINS also interacting with each other in this complex. When the crystal structure of human Cdc45 (at 2.1 Å) was subsequently published [9] the protein was found to harbor twenty α-helices and eleven β-strands, designated α1-20 and β1-11. We identified the PIP box (residues 308 to 314) to localize to the α12 helix (Fig. 4a and 4b upper left panel). Docking of the human Cdc45 crystal structure with the cryo-EM structure of the *Drosophila* CMG complex revealed that the α10-14 bundle was wedged between the Mcm2 and Mcm5 subunits [9]. As the Mcm2/5 interface forms the gate whose opening allows the heterohexamer to encircle the DNA template, the wedging of Cdc45 at this interface ensures this gate remains shut thereafter, allowing the core helicase to move along. The location of the PIP box in this **<u>C</u>**omplex **<u>I</u>**nteraction **<u>D</u>**omain (CID) that interacts with Mcm2/Mcm5 (Fig. 4b upper left panel) limits the possibility of the human Cdc45 PIP box being involved in interactions with PCNA *in vivo*, at least when Cdc45 exists as part of the CMG complex.

**Fig. 4:**
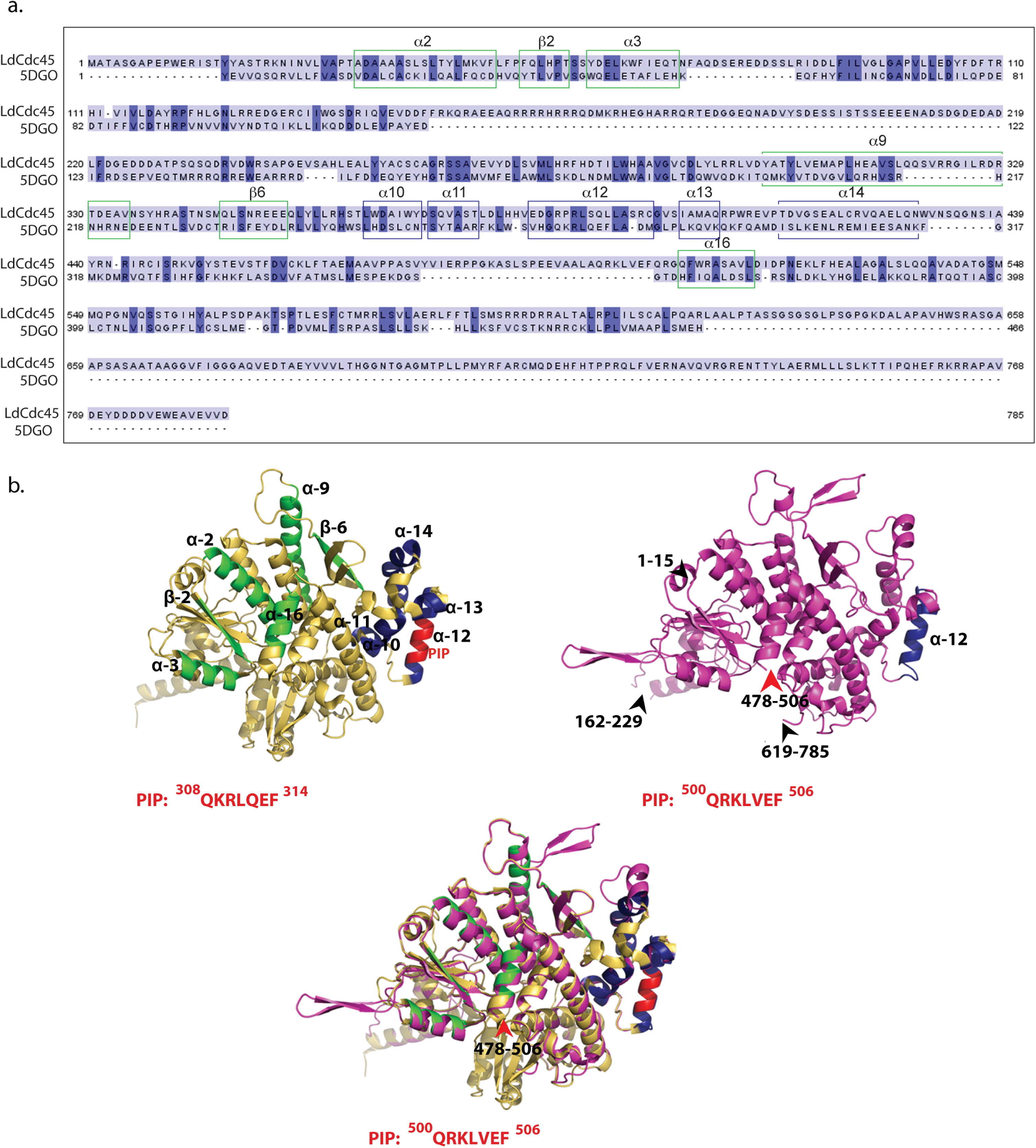
Structural analysis of *Leishmania donovani* Cdc45. **a.** Sequence alignment of *Leishmania donovani* Cdc45 against human Cdc45 generated by Phyre2. Based on structural analysis of human Cdc45 by Simon *et al* [9], α10-α14 are helices that form the Mcm2/5 binding face. All other marked helices and β strands represent regions involved in interaction with GINS **b.** Upper left panel: Human Cdc45 image derived from crystal structure (PDB ID: 5DGO) as reported by Simon *et al* [9]. Navy blue regions: Complex Interaction Domain (CID) that interfaces with Mcm2/5. Green regions: Domains that interface with GINS complex. Red region: PIP box. Sequence of PIP box indicated below structure. Upper right panel: Ribbon representation (magenta) of 3D model of *Leishmania donovani* Cdc45 modelled using Phyre2 against human Cdc45 (PDB ID: 5DGO) as template. Black arrowheads and associated numbers indicate amino acid stretches that have been excluded by Phyre2 during modeling. Red arrowhead and associated numbers indicates location of PIP box (also excluded during modeling). Sequence of PIP box indicated below structure. Navy blue region indicates α12 helix. Lower panel: View of superimposed structures of LdCdc45 and 5DGO, using PyMOL. Though showing overall structural similarity (RMSD value 2.2Å), the PIP motifs of the two structures do not overlap.

To determine the location of the PIP box in LdCdc45 the 3D structure of LdCdc45 was acquired by modeling using Phyre 2.0 ([30]; details in Methods) and the modeled structure was analyzed using PyMOL, an interface that allowed the Cdc45 protein structures to be superimposed to deduce structural alignments. A view of the superimposed models of LdCdc45 and human Cdc45 (5DGO) revealed that while the overall structure of LdCdc45 is similar to that of human Cdc45 a few structural variations exist (Fig. 4b upper right and lower panels). Some regions of LdCdc45 were excluded from the obtained model (marked by arrowheads in Fig. 4b upper right panel). It was observed that while the α12 helix of the CID was conserved in LdCdc45, the PIP box of LdCdc45 did not localize to it. Rather, it was placed on the opposite face of Cdc45 with respect to the CID, suggesting its availability for interactions with PCNA.

### Cdc45 interacts with PCNA in whole cell extracts and in vitro via the PIP box

As the LdCdc45 PIP box appeared to be favourably positioned for interaction with PCNA we examined the possibility of the protein interacting with PCNA in whole cell extracts. Thus, PCNA was immunoprecipitated from cell extracts of asynchronously growing *Leishmania* promastigotes that were expressing Cdc45-FLAG, and the immunoprecipitates were analyzed by western blotting for co-immunoprecipitating Cdc45-FLAG using FLAG antibodies. The data presented in Fig. 5a demonstrated that Cdc45 and PCNA interacted stably in *Leishmania* whole cell extracts.

**Fig. 5:**
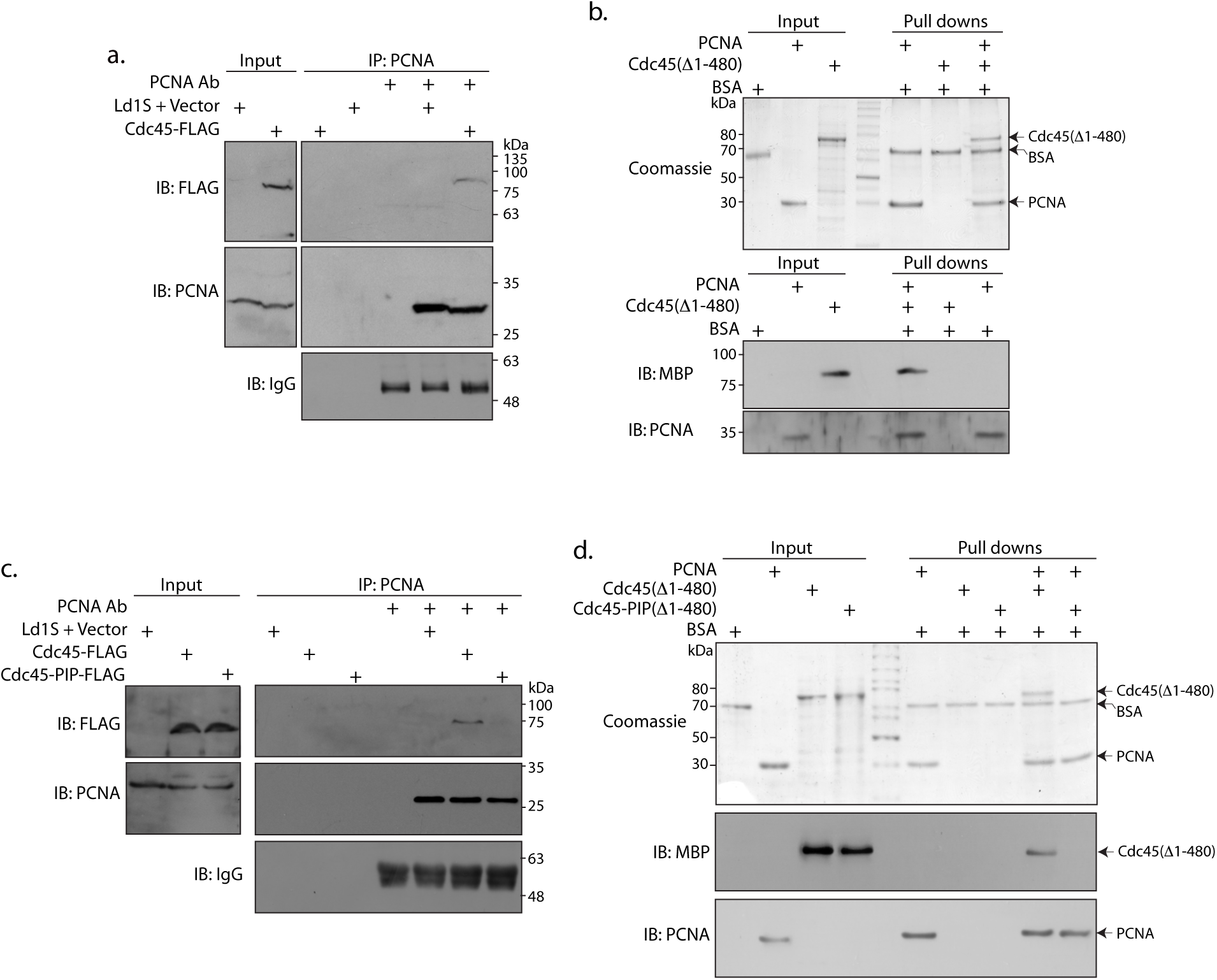
LdCdc45 interacts with PCNA via its PIP box. **a** and **c.** Western blot analysis of PCNA immunoprecipitates from lysates of cells expressing **a.** Cdc45-FLAG **c.** Cdc45-FLAG or Cdc45-PIP-FLAG. Immunoblots were probed with anti-FLAG (Sigma, 1:1000 dil), anti-PCNA (raised earlier in the lab, 1:2500 dil) and anti-IgG (Jackson ImmunoResearch, 1:10000 dil) antibodies. **b.** and **d.** Analysis of pull-down reactions using **b.** PCNA and MBP-Cdc45Δ1-480 **d.** PCNA and MBP-Cdc45Δ1-480 or PCNA and MBP-Cdc45-PIPΔ1-480, by Coomassie staining and western blotting. Immunoblots were probed with anti-MBP (Sigma, 1:12000 dil) and anti-PCNA (1:2500 dil) antibodies.

To ascertain if the Cdc45-PCNA interaction was direct we incubated the two recombinant proteins expressed in *E.coli* in appropriate buffer (described in Methods) and pulled down the His-tagged PCNA using cobalt beads. As evident from Fig. 5b (Coomassie stain as well as western blot analysis using anti-MBP antibodies), the MBP-Cdc45Δ1-480 protein was also pulled down along with PCNA. The role of the PIP box in mediating the Cdc45-PCNA interaction was assessed by creating a PIP box mutant where the **<u>Q</u>**RK**<u>L</u>**VE**<u>F</u>** sequence was altered to **<u>A</u>**RK**<u>A</u>**VE**<u>A</u>**, and expressing the mutant protein in *Leishmania* promastigotes as well as in *E.coli* (Supplementary Methods). Circular dichroism analysis of the recombinant MBP-Cdc45-PIPΔ1-480 protein showed that there were no gross structural differences between the wild type and mutant MBP-Cdc45Δ1-480 proteins (S4a Fig). PCNA immunoprecipitates from whole cell extracts of transfectant cells expressing Cdc45-PIP-FLAG did not carry Cdc45-PIP-FLAG (Fig. 5c), and MBP-Cdc45-PIPΔ1-480 was not pulled down along with PCNA in direct pull-downs between the recombinant proteins (Fig. 5d, Coomassie stain and western blot analysis), signifying the Cdc45-PCNA interaction to be mediated via the PIP box.

To determine if the PIP mutations disrupted the association of Cdc45 with Mcm2-7, LdCdc45-FLAG and LdCdc45-PIP-FLAG immunoprecipitates were analyzed for co-immunoprecipitating Mcm2-7 complex using anti-Mcm4 antibodies already available in the lab [20]. It was observed that Mcm4 co-immunoprecipitated with Cdc45-FLAG and Cdc45-PIP-FLAG to more or less equivalent extent (S4b Fig).

When Simon *et al* [9] docked the human Cdc45 crystal structure in the CMG EM map they identified the GINS-binding surface of Cdc45 to be composite, comprising the α2, α3, α9, α16, β2 and β6 regions. All of these regions were conserved in the LdCdc45 structure we obtained in modeling studies, and the PIP box did not localize to any of these regions (Figure 4a upper right and lower panels). Therefore, the PIP mutations were not expected to negatively impact Cdc45-GINS interactions either. This was experimentally examined by direct pull-downs between the recombinant Cdc45 and Psf1 (GINS subunit interacting with Cdc45) proteins as no antibodies to GINS subunits were available. For this, the *Leishmania donovani psf1* gene was cloned as described in Supplementary Methods, and sequenced (GenBank Accession no. MN612784). The recombinant Psf1 was used in the pull-down reactions, and as seen in western blot analysis, MBP-Cdc45Δ1-480 and MBP-Cdc45-PIPΔ1-480 associated with Psf1 to comparable degrees (S4c Fig). Taken together, these data illustrated that the PIP mutations did not perturb the integrity of the CMG complex.

### Cdc45 PIP box is essential for Leishmania cell survival

The physiological importance of the Cdc45-PCNA interaction and the Cdc45 PIP box *in vivo* was assessed by analyzing the growth and cell cycle patterns of *cdc45*^-/-/+^ parasites expressing Cdc45-PIP-FLAG episomally. Expression of Cdc45-PIP-FLAG was confirmed by western blot analysis (Fig. 6a). It was observed that the ectopic expression of Cdc45-PIP-FLAG did not rescue the growth and cell cycle defects associated with Cdc45 depletion in *cdc45*^-/-/+^ cells (Figs. 6b and 6c, compare *cdc45*^-/-/+^::Cdc45-PIP with *cdc45*^-/-/+^::Cdc45). Furthermore, all attempts to create *cdc45*-nulls using *cdc45*^-/-/+^::Cdc45-PIP cells as the background strain failed, although we were able to create nulls using *cdc45*^-/-/+^::Cdc45 cells as the background strain in parallel. These data underscore the importance of the PIP box of Cdc45 in mediating *Leishmania* cell survival.

**Fig. 6:**
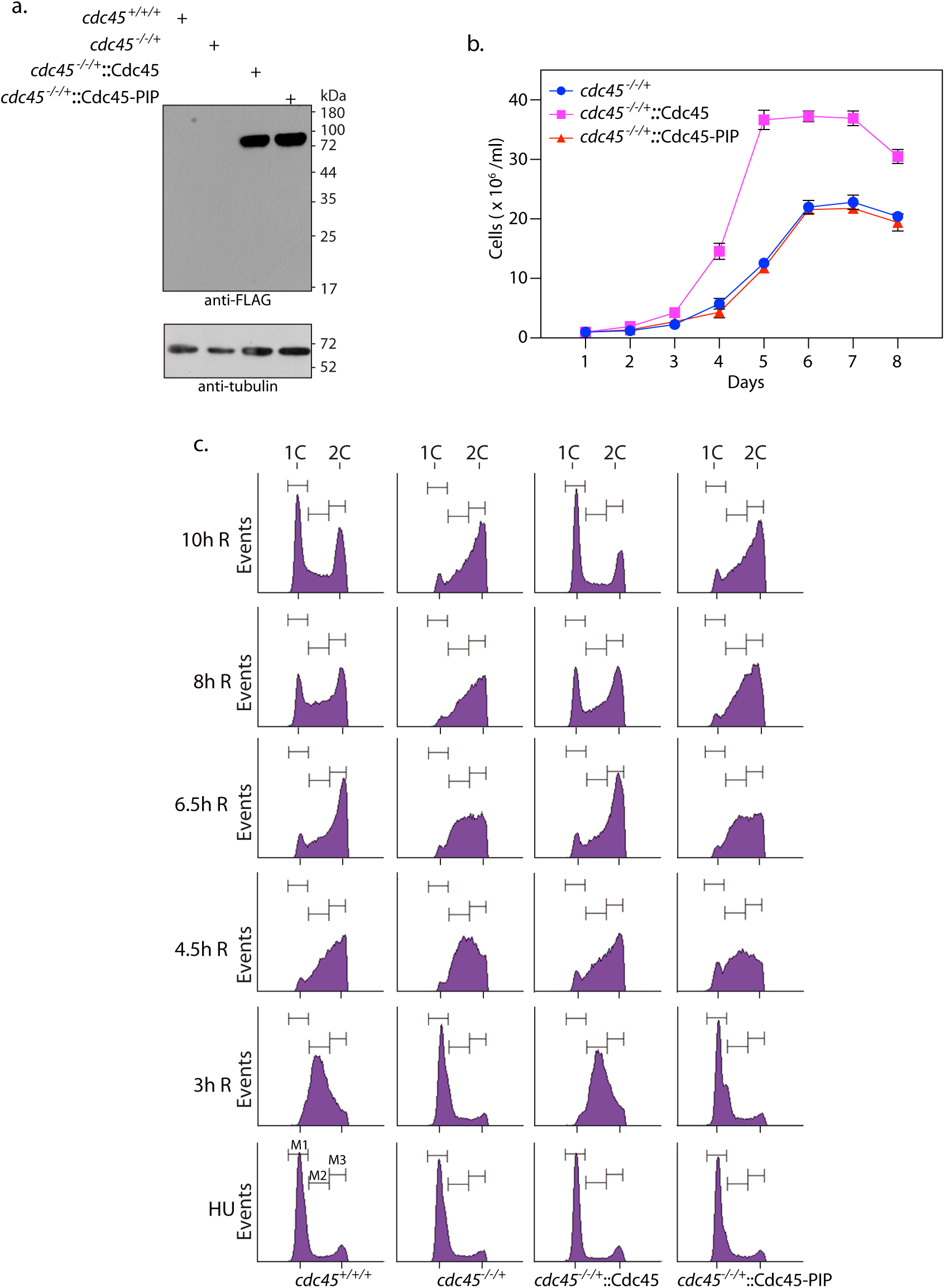
Cdc45-PIP box is essential for *Leishmania* cell survival. **a.** Western blot analysis of whole cell lysates isolated from transfectant *cdc45^-/-/+^* promastigotes expressing Cdc45-FLAG or Cdc45-PIP-FLAG. **b.** Analysis of growth patterns of transfectant *cdc45^-/-/+^*promastigotes expressing Cdc45-FLAG or Cdc45-PIP-FLAG. Graph presents averages of three experiments with error bars indicating standard deviation. **c.** Flow cytometry analysis of transfectant *cdc45^-/-/+^* promastigotes expressing Cdc45-FLAG or Cdc45-PIP-FLAG. The experiment was performed thrice, and the data from one experiment is presented here. 30000 events were analyzed at every sampling. Sampling times are indicated along the y axes and cell types are indicated below the histogram columns.

To examine the *in vivo* impact of the Cdc45-PCNA interaction, chromatin-bound protein fractions of *cdc45*^-/-/+^::Cdc45 and *cdc45*^-/-/+^::Cdc45-PIP cells were analyzed for PCNA binding. The soluble and chromatin-bound protein fractions isolated from logarithmically growing promastigotes (described in Methods) were analyzed for the quality of chromatin fractionation by western blot analysis using anti-H4acetylK4 antibodies [31], and the same blots were then probed for PCNA. The data presented in Figs. 7a and 7d revealed that the amount of chromatin-bound PCNA in *cdc45*^-/-/+^::Cdc45-PIP cells was approximately half that detected in *cdc45*^-/-/+^::Cdc45 cells. The amount of chromatin-associated PCNA in promastigotes that had been incubated in hydroxyurea for 8 hours was comparable between both parasite types (Figs. 7b and 7d), in keeping with the fact that the cells were non-replicating at this stage. Upon examining chromatin-bound protein fractions of cells two hours after release from hydroxyurea-induced block we observed that while the amount of chromatin-bound PCNA was significantly higher in both cell types (in keeping with the fact that both cell types had now entered S phase, with *cdc45*^-/-/+^::Cdc45 cells advancing more than *cdc45*^-/-/+^::Cdc45-PIP cells), the amount of chromatin-bound PCNA was significantly higher in *cdc45*^-/-/+^::Cdc45 cells (Fig. 7c and 7d). These data suggest that the Cdc45-PCNA interaction that is mediated by the PIP motif of Cdc45 may play a role in recruiting PCNA-polymerase complexes to the advancing replication fork, or may stabilize the association of PCNA-polymerase complexes with template DNA during active DNA replication. The amount of chromatin-bound Cdc45 and Mcm2-7 (determined using Mcm4 as marker) was not affected by the PIP mutations (Fig. 7c), suggesting that CMG complex loading on template DNA was unaffected.

**Fig. 7:**
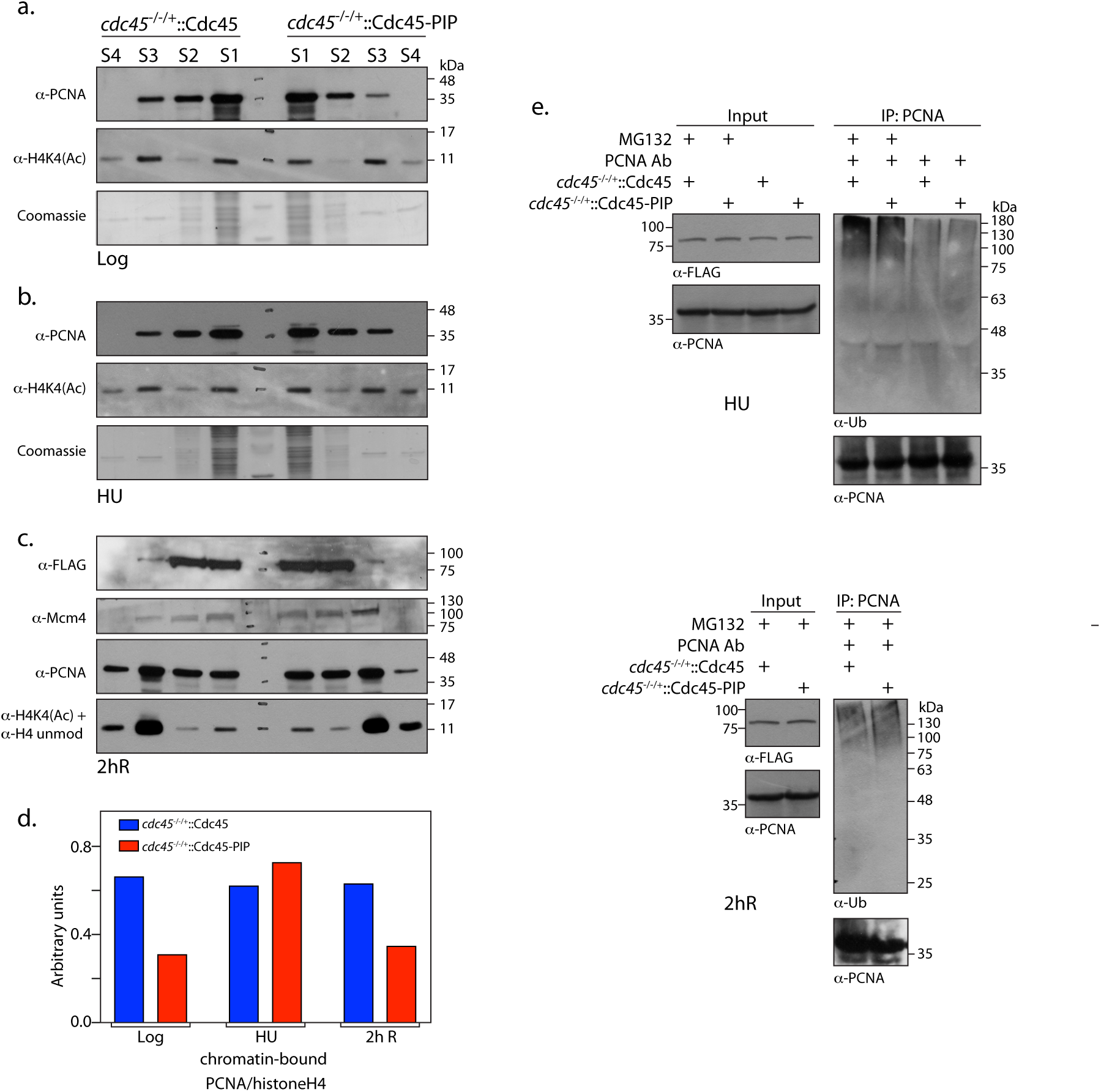
Cdc45 helps recruit/stabilize PCNA-polymerase complexes on chromatin during active DNA replication. Analysis of soluble and chromatin-bound protein fractions isolated from *cdc45*^-/-/+^::Cdc45 and *cdc45*^-/-/+^::Cdc45-PIP cells. Each experiment was carried out twice. One data set for each is presented here. Reactions resolved on 12% SDS-PAGE were probed with various antibodies as indicated. S1, S2 – soluble fractions; S3, S4 – DNA-associated fractions. **a.** isolated from 5 ×10^7^ logarithmically growing cells. **b.** isolated from 5 × 10^7^ HU treated cells **c.** isolated from 2 ×10^9^ synchronized cells 2 hours after release from block. **d.** ratio of chromatin-bound PCNA (S3+S4): histone H4 (S3+S4) was determined at different time points (log, HU, 2hR) by quantification using ImageJ software. **e.** Analysis of PCNA immunoprecipitates from whole cell extracts of *cdc45*^-/-/+^::Cdc45 and *cdc45*^-/-/+^::Cdc45-PIP cells. Cells were treated with 5 mM HU for 8 hours, with 20 µM MG132 being added 5 hours into the treatment (upper half of figure). In case of cells that were harvested 2 hours after release from HU-induced block, 20 µM MG132 was also added to the cells at the time of release from HU (lower half of figure). The immunoprecipitates were resolved on SDS-PAGE and analysed for ubiquitinated PCNA using anti-Ub antibodies (Santa Cruz, 1:1000 dil). The blot was subsequently probed with anti-PCNA antibody (1:2500 dil).

In view of the data in Figs. 7a-7d we examined the possibility of chromatin-bound PCNA being degraded more rapidly in absence of interaction with Cdc45. Considering that most intracellular proteins are degraded by the ubiquitin proteasome pathway (UPP) where proteins destined for degradation are marked by covalent tagging with ubiquitin, we examined PCNA immunoprecipitates of extracts isolated from *cdc45*^-/-/+^::Cdc45 and *cdc45*^-/-/+^::Cdc45-PIP cells to see if PCNA was differentially tagged with poly-Ub in these two cell types. For this, HU-synchronized cells were incubated with MG132 (a peptide aldehyde that inhibits serine and cysteine proteases of the 26S proteasome; 20 µM), which was added to the culture 3 hours before release from hydroxyurea-induced block as well as at the time of release from block. Cell lysates were isolated after 8 hours in hydroxyurea as well as 2 hours after release from HU-induced block. When PCNA immunoprecipitates from these cell extracts were analyzed for ubiquitinated PCNA using anti-Ub antibodies (Santa Cruz Biotechnologies, a kind gift from Dr. Alo Nag and Dr. Sagar Sengupta), it was observed that the extent of PCNA polyubiquitination was comparable in immunoprecipitates of *cdc45*^-/-/+^::Cdc45 and *cdc45*^-/-/+^::Cdc45-PIP (Fig. 7e), at both time points examined. This suggests that the Cdc45-PCNA interaction does not modulate PCNA degradation. Considering the data from Figs. 7a-7e, we conclude that the Cdc45-PCNA interaction either helps recruit PCNA-polymerase complexes to template DNA, or stabilizes the interaction of PCNA-polymerase complexes with template DNA, without playing any direct role in PCNA degradation.

### Cdc45 PIP box is essential for survival of Schizosaccharomyces pombe

As indicated in S3 Fig, *Drosophila* and *S. pombe* Cdc45 proteins also carry PIP boxes. The location of the PIP box of *Drosophila* Cdc45 in the α20 helix near the C-terminus of the protein, away from the <u>C</u>omplex <u>I</u>nteraction <u>D</u>omain (CID) which is responsible for the interaction of Cdc45 with the Mcm2/5 subunits, supports the possibility of interactions with PCNA (S5 Fig; *Drosophila* Cdc45 image derived from the electron microscopy structure, PDB ID: 6RAW [32]). When the *S. pombe* Cdc45 was modeled against the human Cdc45 crystal structure the PIP box of *S. pombe* Cdc45 was found to lie away from the CID, in the extension of the α16 helix (Fig. 8a). Considering this fact, we investigated the importance of the Cdc45 PIP motif in *S. pombe* by carrying out complementation assays using the *S. pombe sna41^goa1^* strain, a kind gift from Prof. Hisao Masai. This strain is a *cdc45 ^ts^*mutant that grows at 25°C, but not at 37°C (unlike wild type *S. pombe* which thrives at 37°C; [33, 34]).

**Fig. 8:**
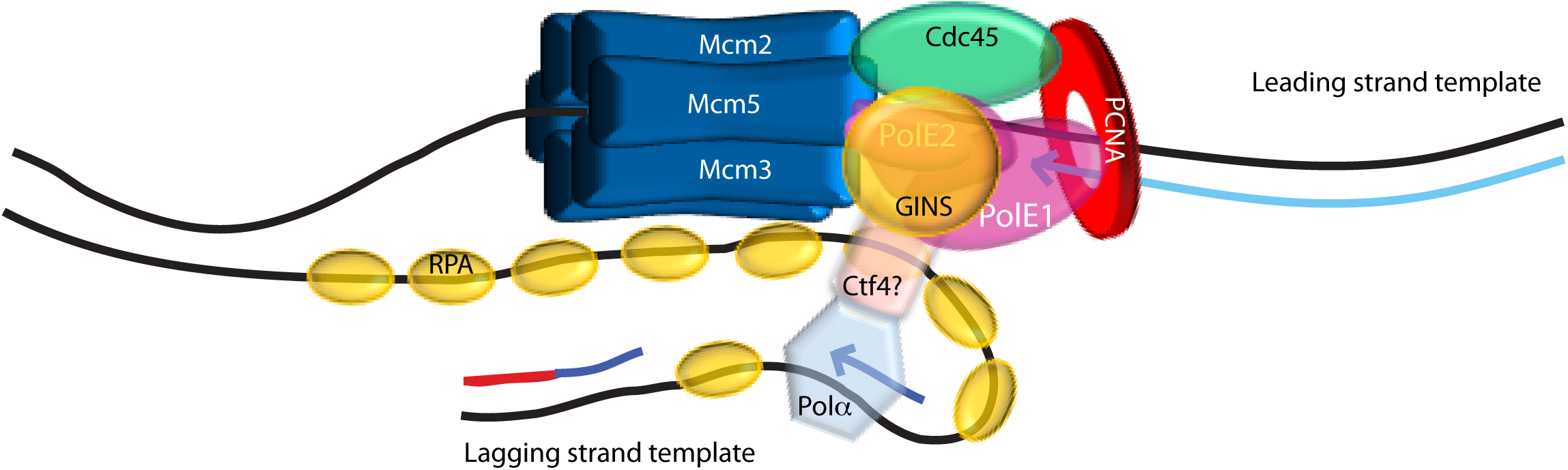
Cdc45-PIP box is essential for *S. pombe* cell survival: **a.** Upper panel: Sequence alignment of *Schizosaccharomyces pombe* Cdc45 against human Cdc45 generated by Phyre2. Lower left panel: Human Cdc45 image derived from crystal structure (PDB ID: 5DGO; [9]). Navy blue region indicates α12 helix, red region depicts PIP box, sequence of PIP box indicated below structure. Lower right panel: Ribbon representation (pink) of 3D model of *Schizosaccharomyces pombe* Cdc45 modelled with Phyre2 using human Cdc45 (PDB ID: 5DGO) as template. Black arrowheads and associated numbers indicate amino acid stretches that have been excluded by Phyre2 during modeling. Navy blue region indicates α12 helix. Red region indicates PIP box. Sequence of PIP box indicated below structure. **b.** Western blot analysis of whole cell lysates isolated from *sna1^goa1^* transformants expressing LdCdc45 (wild type or PIP mutant) proteins. Two clones of each type were analyzed, using anti-FLAG antibodies (1:1000 dil). **c.** Complementation assays. *sna1^goa1^*transformants expressing LdCdc45 (wild type or PIP mutant) proteins were streaked on EMM2 plates (with necessary supplements) and incubated at permissive and non-permissive temperatures to assess functional complementation. Transformant expressing SpCdc45 served as positive control for complementation. **d.** Western blot analysis of whole cell lysates isolated from *sna1^goa1^* transformants expressing SpCdc45 (non-tagged wild type or His-tagged PIP mutant) proteins. Two clones of SpCdc45-PIP were analyzed, using anti-His antibodies (1:10000 dil). **e.** Complementation assays. *sna1^goa1^* transformants expressing SpCdc45 (wild type or PIP mutant) proteins were streaked on EMM2 plates and incubated at permissive and non-permissive temperatures.

To carry out complementation experiments the *Leishmania* Cdc45-FLAG (wild type and PIP mutant) and SpCdc45 (wild type, non-tagged) proteins were expressed in *sna41^goa1^* at 25°C (described in Supplementary Methods; Fig. 8b shows western blot analysis of expression of Cdc45-FLAG and Cdc45-PIP-FLAG in two clones of each type). The ability of the *Leishmania* Cdc45-FLAG proteins to overcome the growth defects of the *sna1^goa1^* cells was examined by streaking out the clones on EMM2 agar plates with necessary supplements, and incubating the plates at 25°C, 30°C and 37°C. It was observed that while SpCdc45 was able to rescue the growth defects of *sna1^goa1^* cells at 37°C, neither LdCdc45-FLAG nor LdCdc45-PIP-FLAG could do so (Fig. 8c). While it is unclear why the wild type LdCdc45-FLAG is unable to complement the *cdc45^ts^* mutation in *sna1^goa1^* cells, it is possibly linked to the fact that the *Leishmania* GINS subunits have a somewhat different structure from those of *S. pombe.* Therefore, LdCdc45 may not be able to interact with *S. pombe* GINS in a productive manner, and an active CMG complex involving LdCdc45 may not form in *S. pombe*.

Hence, to analyze the importance of the Cdc45-PIP motif in *S. pombe* we created an *S. pombe* Cdc45-PIP mutant protein. The SpCdc45 PIP sequence QEWLHNFY was mutated to AEWAHNAY and the mutant protein expressed with a C-terminal His tag in *sna1^goa1^*cells (western blot analysis of SpCdc45-PIP seen in Fig. 8d). When complementation analysis was carried out as earlier at 25°C, 30°C and 37°C, it was found that while wild type SpCdc45 complemented the inherent growth defects of *sna1^goa1^*the SpCdc45-PIP mutant protein did not (Fig. 8e). These data lead us to conclude that the Cdc45 PIP motif is essential not only for survival and propagation of *Leishmania* but for the survival and propagation of *S. pombe* as well.

## Discussion

Cdc45 (cell division cycle 45), originally identified in genetic screens for yeast mutants defective in cell cycle progression [35], is now well established as a protein required for DNA replication initiation as well as elongation of the DNA chains being synthesized. As part of a complex with the heterohexameric MCM and heterotetrameric GINS, it forms the replicative helicase that advances with the replication fork: the CMG complex. The association of Cdc45 and GINS with Mcm2-7 is stabilized by Mcm10, which has also been demonstrated to be essential for the activation of CMG helicase activity [4, 36]. Cdc45 is also involved in actively loading RPA, the ssDNA-binding protein, onto the newly unwound DNA, thus allowing further association of the enzymatic machinery required for replication initiation and elongation [37]. Using a tethered bead assay, the *Saccharomyces cerevisiae* leading strand replisome has been visualized at single molecule level and found to comprise twenty-four proteins including the eleven-subunit CMG complex, DNA pol ε, PCNA, RFC clamp loader and RPA [38]. As DNA replication commences, the replisome complexes advance along with the replication forks bidirectionally, with the CMG unwinding the DNA ahead using energy derived from ATP hydrolysis, to enable template-dependent DNA synthesis (reviewed in [2]). The role of Cdc45 is conserved across eukaryotes, and the data presented in Figs. 1-3 demonstrates that it plays a role in DNA replication in *Leishmania donovani* as well.

Proliferating cell nuclear antigen is a highly conserved homotrimeric protein that itself does not possess any enzymatic activity, yet regulates a multitude of cellular processes involving DNA metabolism by virtue of its ability to interact with a vast number of proteins, both in concerted and sequential manner. Among the repertoire of its direct partners are included several components of the DNA replication machinery: replication factor C (RFC), DNA pol α, DNA polymerase δ, DNA polymerase ε, the flap endonuclease FEN1, DNA ligase I (reviewed in [29, 39]). The front face of the “sliding clamp” forms the interface for protein-protein interactions. A large number of the proteins which directly interact with PCNA are typified by the presence of a conserved sequence called the PCNA-interacting-peptide (PIP) motif through which they bind to PCNA, by the insertion of the PIP box into a hydrophobic pocket beneath the flexible interdomain connecting loop (IDCL) of PCNA (the region that connects the two globular domains of each PCNA monomer). As each trimer has three such pockets, it allows for the simultaneous binding of three different partner proteins. The identification of a PIP box in *Leishmania* Cdc45 led us to investigate the possibility of Cdc45 being yet another of the many partners of PCNA.

Structural analysis of LdCdc45 revealed to us that the PIP box lay opposite to the Cdc45/Mcm2-7 and Cdc45/GINS interfaces (Fig. 4), supporting the possibility of interaction with PCNA. The data presented in Fig. 5 confirmed that Cdc45 and PCNA interacted with each other not only in whole cell extracts, but directly *in vitro* also, and furthermore, this interaction occurred through the Cdc45 PIP box. Our laboratory has previously shown *Leishmania* Mcm4 to interact with PCNA in whole cell extracts, however, no evidence of a direct interaction was found [20]. Based on the data in Figs. 5 and S4b, it appears that the interaction of Mcm4 with PCNA is in fact through Cdc45. While a PIP motif was identified in LdMcm4 [20], subsequent alignment with the later published ScMcm2-7 crystal structure [40] revealed the LdMcm4 PIP box to lie in the zinc finger domain that is located at the inter-hexamer interface, signifying its involvement in MCM double hexamer formation.

In pursuit of the possible *in vivo* role of the Cdc45-PCNA interaction we took into consideration previous biochemical and structural studies (including single particle electron cryo-microscopy studies) which have revealed that DNA pol ε (the leading strand DNA polymerase) interacts directly with the CMG complex as it moves along the leading strand. Studies from budding yeast have found the interaction of DNA pol ε with CMG to occur primarily via the interaction of the non-catalytic Dpb2 subunit (PolE2 in humans) with Psf1 (of GINS) and Mcm5, and via the interaction of the non-catalytic C-terminal domain of the Pol2 subunit (PolE1 in humans) with Mcm5 and Cdc45 [11,12,41–44]. While no direct interaction between CMG and DNA pol δ (the lagging strand polymerase) has been reported, CMG interacts with Pol α on the lagging strand indirectly through the adaptor protein Ctf4 (chromosome transmission fidelity; first identified in yeast screens, [10, 12]. Using *in vitro* replication assays with yeast proteins Yeeles *et al* [45] showed that while DNA polymerase δ was important for initiating leading strand replication, replication elongation was efficiently continued by DNA polymerase ε. Secondly, PCNA was critical to maximal rates of leading strand DNA synthesis by DNA polymerase ε. The authors proposed that while the CMG - Pol ε interaction may be responsible for hitching Pol ε to the advancing replication fork, the PCNA –Pol ε interaction increased the processivity of the polymerase during DNA synthesis *per se,* thus increasing the replication rate considerably. Thus, the unwinding of the DNA duplex by CMG remains tightly coupled to the synthesis of DNA by Pol ε. Based on experimental results presented in Figure 7 of this manuscript we hypothesize that the Cdc45-PCNA interaction may further stabilize the interaction of DNA pol ε with CMG, thus anchoring the polymerase more firmly with the advancing leading strand replisome and maximizing efficiency of DNA synthesis (Fig. 9).

**Fig. 9:**
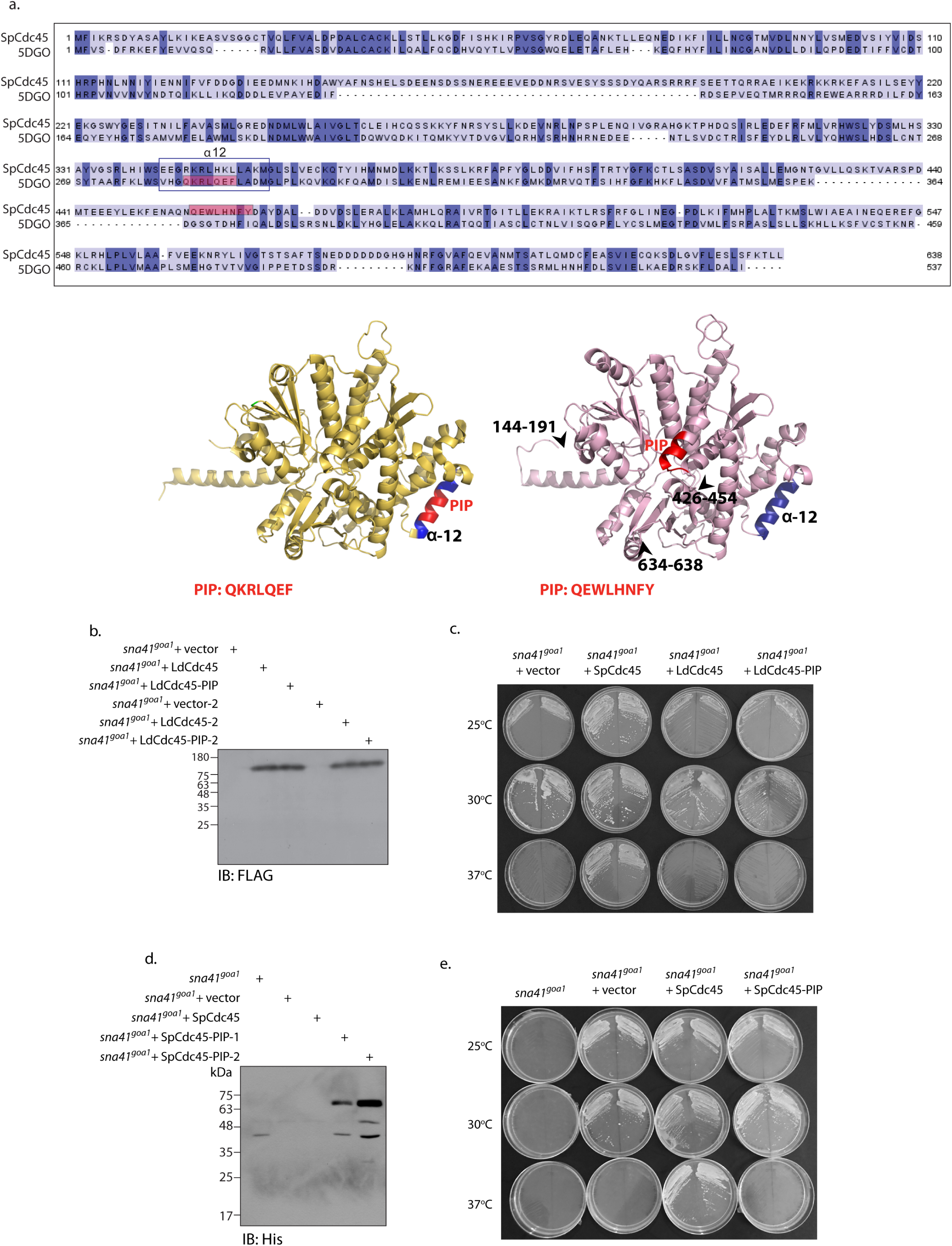
The role of Cdc45-PCNA interaction at the replication fork. The active replisome on the leading strand template includes several proteins such as the Cdc45-MCM-GINS complex, DNA pol ε, and PCNA. Here, the heterotetrameric GINS is represented as a single entity. DNA pol ε is depicted with only two subunits: PolE1 (bi-lobed) and PolE2, as the other two subunits have not been identified in trypanosomatids. No Ctf4 has yet been identified in trypanosomatids. The interaction of Cdc45 with PCNA, the sliding clamp processivity factor of the replicative polymerases, may further stabilize the polymerase on the advancing replisome machinery, thus enhancing the efficiency of replication.

The Cdc45 PIP box was found to be essential for *Leishmania* cell survival, as the Cdc45-PIP mutant protein did not rescue the severe defects associated with Cdc45 depletion (Fig. 6) and ectopic expression of Cdc45-PIP protein did not support knockout of the third genomic allele in *cdc45^-/-/+^* cells. Complementation assays revealed that the *S. pombe* Cdc45 PIP box was also essential for cell survival (Fig. 8), implicating the importance of a PIP-mediated Cdc45-PCNA interaction in *S. pombe* also. Separate detailed investigations are necessary to ascertain if Cdc45 and PCNA interact in *S. pombe* and *Drosophila*. The findings presented in this paper underscore the fact that there are diverse facets in modes of DNA replication among eukaryotes even though the process is broadly conserved across them, also pointing to the relevance of studying this process in non-conventional eukaryotes like *Leishmania*.

## Materials and Methods

### Leishmania cultures and manipulations

*Leishmania donovani* 1S promastigotes were cultured as earlier [22] in medium M199 (Lonza, Switzerland) supplemented with fetal bovine serum (10%; Invitrogen), hemin and adenine (Sigma Aldrich, USA). Isolation of whole cell extracts, analysis of growth patterns and generation time, synchronization regimes and flow cytometry analysis methods, transfections of *Leishmania* promastigotes, and creation and maintenance of clonal lines were done as described in Supplementary Methods (Supporting Information).

### Schizosaccharomyces pombe cultures and manipulations

*S. pombe* cultures (wild type or *sna41^goa1^* mutant strain; kind gifts from Dr. Nimisha Sharma and Prof. Hisao Masai respectively) were grown and maintained at 25°C on solid YES medium, or EMM2 supplemented with adenine, uracil, leucine, lysine and histidine (225 mg/l each). *S. pombe* transformations and complementation assays were done as described in Supplementary Methods (Supporting Information).

### Real time PCR analysis

The PureLink RNA mini kit (Invitrogen, USA), iScript cDNA synthesis kit and iTaq Universal SYBR green supermix (both from Bio-Rad Laboratories, USA) were used for isolation of total RNA, cDNA synthesis and real time PCR analyses respectively. *cdc45* expression was analyzed by primers Cdc45-RT-F and Cdc45-RT-R (5’-CTGCGCCTTCAGCGTCTG-3’ and 5’-TGTAGTCTTCAACGACAGG-3’ respectively), and tubulin expression was analyzed by primers Tub-RT-F1 and Tub-RT-R2 (5’-CTTCAAGTGCGGCATCAACTA-3’ and 5’-TTAGTACTCCTCGACGTCCTC-3’ respectively). The reactions were run in the CFX96 Real Time System (Bio-Rad Laboratories, USA) and *cdc45* expression analyzed in knockout cells in relation to wild type cells using the 2^-ΔΔC^_T_ method as earlier [46]. The experiment was done three times, with reactions being set up in triplicate in each experiment. Bars represent average values of the three experiments. Error bars represent standard deviation. Student’s t-test (two-tailed) was applied to determine the significance of the data.

### EdU labeling analysis

EdU labeling analysis was performed as described [46]. Briefly, promastigotes were incubated in 5 mM hydroxyurea for 8 hours, released into drug-free M199, and cell aliquots removed at different time intervals thereafter for 15 minute pulses with 5-ethynyl-2-deoxyuridine (EdU). EdU uptake was detected with the help of the Click-iT EdU Imaging Kit (Invitrogen) followed by imaging of cells using a LeicaTCS SP5 confocal microscope with a 100X objective.

### Macrophage infection experiment

J774A.1 cells were infected with *Leishmania* metacyclics as described [46]. Three biological replicates of the experiment were set up and the mean values of the three experiments are presented in a bar chart, with standard deviation being depicted by error bars. The significance of the data obtained was analyzed by applying Student’s t-test (two-tailed).

### Homology modelling and structural analysis

The structures of *Leishmania* and *Schizosaccharomyces pombe* Cdc45 were built using Phyre2 via homology modeling in normal mode (http://www.sbg.bio.ic.ac.uk/phyre2) and were visualized and further analyzed using PyMOL (http://pymol.org). In both cases, the two most reliable 3D models that were obtained were against *Saccharomyces cerevisiae* Cdc45 (PDB ID: 3JC6 [47]) and human Cdc45 (PDB ID: 5DGO [9]). As the coverage and percent identity were in similar range (*Leishmania* Cdc45: 82% coverage with 25% identity against 3JC6, 77% coverage with 23% identity against 5DGO; *Schizosaccharomyces pombe* Cdc45: 99% coverage with 38% identity against 3JC6, 99% coverage with 32% identity against 5DGO), and the 5DGO was a crystal structure determined at a higher resolution than the 3JC6 electron microscopy structure (2.1 Å versus 3.7 Å), the 3D models obtained against human Cdc45 was considered for further analysis. An additional factor that was considered was that human Cdc45 has a PIP motif, while ScCdc45 does not.

### Chromatin binding analyses

Isolation of soluble and DNA-associated protein fractions from promastigotes was carried out as detailed earlier [31]. The fractions (S1, S2: soluble fractions; S3, S4: DNA-associated fractions) were analyzed by probing them in western blots.

### Immunoprecipitations and pulldowns

Whole cell lysates were isolated from logarithmically growing transfectant *Leishmania* promastigotes (2 × 10^9^ cells) expressing either Cdc45-FLAG or Cdc45-PIP-FLAG from plasmids pXG/Cdc45-FLAG and pXG/Cdc45-PIP-FLAG respectively, and two-thirds of the isolated lysates were used for immunoprecipitation analysis. For PCNA immunoprecipitations, the anti-PCNA antibodies (10 µl; raised in rabbit previously in the lab [21]) were first bound to protein A sepharose beads (40 µl protein A sepharose/CL6B sepharose 1:1 slurry mix) by incubation on ice for one hour with periodic mixing. This was followed by washing of the beads with 1XPBS-0.1%TX-100 to remove unbound antibody fraction, and adding the cell lysates that had been treated with 50U of DNaseI. Immunoprecipitation was allowed to proceed overnight at 4°C, followed by washing the beads extensively before adding SDS sample buffer and boiling. The entire mix was analyzed in western blotting. Antibodies used to probe PCNA immunoprecipitates included anti-FLAG (mouse monoclonal, Sigma, 1:1000 dil), anti-PCNA (rabbit; 1:2500 dil), anti-Ub (mouse monoclonal, Santa Cruz Biotechnology, 1:1000) and anti-IgG (rabbit, Jackson Laboratories, 1:10000 dil) antibodies. For immunoprecipitating FLAG-tagged proteins, the cell lysates were incubated with M2-agarose beads (Sigma; 40 µl M2-agarose /CL6B sepharose 1:1 slurry mix) overnight at 4°C, before washing and analysis.

For direct pull-downs using PCNA, recombinant His-tagged PCNA was purified as earlier [21]. Expression of the recombinant MBP-Cdc45Δ1-480 proteins (wild type and PIP mutant) was induced at 37°C (using IPTG) in BL21 CodonPlus cells harbouring plasmids pMAL/Cdc45Δ1-480 or pMAL/Cdc45-PIPΔ1-480. To carry out the pull-down, 25 pmoles of purified PCNA (trimeric protein) were incubated with 130-145 µg of bacterial cell extracts containing the overexpressed MBP-Cdc45Δ1-480 protein (wild type or mutant), in 1X PBS (total mix volume 500 µl) for 2 h at 4°C with gentle mixing using a nutator mixer. 50 µl cobalt beads (Talon metal affinity beads; BD Bioscience) that had been pre-blocked with BSA for two hours (by incubation of beads in 400 µl 1X PBS containing 100 µg BSA, followed by two washes to remove excess BSA) were then added to the pull-down reaction mix. The reaction was further incubated at 4°C for 30 min with mixing. After removing the unbound fraction by low speed centrifugation the beads were washed with 100 mM Tris.Cl (pH 8), 750 mM NaCl and the bound proteins eluted using 100 mM Tris.Cl (pH 8), 300 mM NaCl, 250 mM imidazole. One-fifth of the elute fractions were analyzed by Coomassie staining and one-hundredth were analyzed by western blotting. Direct pull-down assays using Psf1 were similarly carried out after purifying the Strep-tagged Psf1 by Strep-Tactin II chromatography and using Talon metal affinity beads for the pull-downs.

### Data availability

GenBank Accession numbers: MN612783 for Ld1S Cdc45; MN612784 for Ld1S Psf1.

## Acknowledgements

We thank Prof. Hisao Masai of the Tokyo Metropolitan Institute of Medical Science for kindly providing us with the *sna1^goa1^*strain. We thank Dr. Nimisha Sharma of the Guru Gobind Singh Indraprastha University, Delhi, for providing us *S. pombe* cultures and pART1 vector. We thank Dr. Aruna Naorem for the pLP-BLP vector. We thank Dr. Sagar Sengupta and Prof. Alo Nag for kindly providing us anti-Ub antibodies. We thank Dr. Vinay Nandicoori for allowing us the use of his laboratory facilities. DNA sequencing, Flow cytometry analyses, and Confocal microscopy were done at the Central Instrumentation Facility, University of Delhi South Campus.

## Supporting Information Legends

### Supporting Information

Supporting Information carries Supplementary Methods describing manipulations of Leishmania promastigotes, manipulations using *S. pombe*, cloning of *Leishmania donovani cdc45* and *psf1* genes, cloning of *S. pombe cdc45* gene, tagging of *cdc45* with *eGFP*, creating *cdc45* knockout and rescue lines, immunofluorescence analysis, and CD spectroscopic analysis.

### Supporting Figures

**S1 Fig: Comparative analysis of Leishmania donovani Cdc45 sequence with Cdc45 of other trypanosomatids**. Clustal Omega analysis of LdCdc45 with Cdc45 of other trypanosomatids viewed using Jalview multiple alignment editor [7]. Black rectangles mark PIP boxes. Colours are indicative of the physico-chemical properties of the amino acids. Pink, aliphatic/hydrophobic; orange/ochre, aromatic; purple, glycine/proline; dark blue, basic; green, hydrophilic; red, acidic; yellow, cysteine.

**S2 Fig: Cdc45 is constitutively nuclear in *Leishmani donovani*. a**. Tagging one *cdc45* genomic allele with *eGFP*. Primers used in screening indicated by arrows. Agarose gels depict screening across replacement junctions, with primer pairs marked below. Lanes 1 – Ld1S, lanes 2 – replacement line. **b.** Western blot analysis of whole cell lysates probed with anti-eGFP antibodies (already available in the lab). Lane 1 – Ld1S, lane 2 – replacement line. **c.** Immunofluorescence analysis of Cdc45 at different cell cycle stages using kinetoplast morphology and number as cell cycle stage marker. G1/early S: one nucleus, one short or roundish kinetoplast; late S/early G2/M: one nucleus, one elongated kinetoplast; G2/M: two nuclei, one kinetoplast; post-mitosis: two nuclei, two kinetoplasts.

**S3 Fig: Comparative analysis of *Leishmania donovani* Cdc45 sequence with Cdc45 of other eukaryotic organisms**. Clustal Omega analysis viewed using Jalview multiple alignment editor [7]. Black rectangles mark PIP boxes. Colours indicative of physico-chemical properties of the residues. Pink, aliphatic/hydrophobic; orange/ochre, aromatic; purple, glycine/proline; dark blue, basic; green, hydrophilic; red, acidic; yellow, cysteine.

**S4 Fig: The PIP mutations do not affect Cdc45-MCM or Cdc45-GINS interactions**. **a**. CD spectra of MBP-Cdc45Δ1-480 and MBP-Cdc45-PIPΔ1-480 are depicted as a measure of mean residue ellipticity. **b**. Analysis of Cdc45-FLAG and Cdc45-PIP-FLAG immunoprecipitates from lysates isolated from transfectant *cdc45*^-/-/+^ cells. Western blot analysis done using anti-Mcm4 antibodies (previously raised in the lab [5], 1:500) and anti-FLAG antibodies (Sigma, 1:1000). c. Analysis of pull-down reaction between MBP-Cdc45Δ1-480 and LdPsf1, and MBP-Cdc45-PIPΔ1-480 and LdPsf1. Western blot analysis done using anti-MBP (Sigma, 1:12000) and anti-His (Sigma, 1:5000).

**S5 Fig: Examining Drosophila Cdc45 for PIP box**. Left panel: Image of human Cdc45 derived from crystal structure PDB ID: 5DGO [8]. Navy blue region: α12 helix. Red region: PIP box, sequence below structure. Right panel: Image of *Drosophila* Cdc45 derived from electron microscopy structure PDB ID: 6RAW [9]. Navy blue region: α12 helix. Red region: PIP box, sequence below structure.

